# Cytosolic acetyl-CoA synthetase (ACSS2) does not generate butyryl- and crotonyl-CoA

**DOI:** 10.1101/2023.12.21.572516

**Authors:** Nour Zeaiter, Laura Belot, Valérie Cunin, Roland Abi Nahed, Malgorzata Tokarska-Schlattner, Audrey Le Gouellec, Carlo Petosa, Saadi Khochbin, Uwe Schlattner

## Abstract

Acetyl and other acyl groups from different short-chain fatty acids (SCFA) competitively modify histones at various lysine sites. To fully understand the functional significance of such histone acylation, a key epigenetic mechanism, it is crucial to characterize the cellular sources of the corresponding acyl-CoA molecules required for the lysine modification. Like acetate, SCFAs such as propionate, butyrate and crotonate are thought to be the substrates used to generate the corresponding acyl-CoAs by enzymes known as acyl-CoA synthetases. The acetyl-CoA synthetase, ACSS2, which produces acetyl-CoA from acetate in the nucleocytoplasmic compartment, has been proposed to also mediate the synthesis of acyl-CoAs such as butyryl- and crotonyl-CoA from the corresponding SCFAs. This idea is now widely accepted and is sparking new research projects. However, based on our direct *in vitro* experiments with purified or recombinant enzymes and structural considerations, we demonstrate that ACSS2 is unable to mediate the generation of non-acetyl acyl-CoAs like butyryl- and crotonyl-CoA. It is therefore essential to re-examine published data and corresponding discussions in the light of this new finding.

## Introduction

Acetyl-CoA synthetase 2 (ACSS2) catalyzes the synthesis of acetyl-CoA from intracellular acetate. Together with ATP citrate lyase, ACSS2 provides a nucleocytosolic source of acetyl-CoA, which is competitively consumed in histone/protein lysine acetylation (Kac), *de novo* lipogenesis and the mevalonate pathway. ACSS2 thus plays key roles for epigenetic regulation and metabolic homeostasis in different pathophysiological settings (e.g. [1, 2]) and has emerged as a new drug target for cancer [3].

Following the pioneering work of Yingming Zhao since 2007, in addition to acetylation, a series of histone lysine acylations was identified, including propionylation (Kpr) and butyrylation (Kbu) [4], crotonylation (Kcr) [5], 2-hydroxyisobutyrylation (2hib) [6], β-hydroxybutyrylation (Kbhb) [7], malonylation (Kmal), succinylation (Ksucc) [8], and others. *In vitro* and *in vivo* experimental data suggest that various histone acetyltransferases (HATs), such as CBP/p300 [9-11], are responsible for these histone acylations. These experimental data imply that the CoA derivatives for all these acyl groups exist in the cell and are available in the nucleus as acyl group donors for HAT-mediated histone modification. Indeed, since all known HATs use acetyl-CoA as the acetyl group donor, other HAT-mediated histone acylations should also depend on a given acyl-CoA, and HATs should be able to accommodate these longer-chain acyl groups in their catalytic site [9].

Given all these data and observations, an important question arises: what are the sources of the different acyl-CoAs used by cellular HATs to carry out these histone acylations? The acyl-CoAs corresponding to several well-studied histone modifications are generated in mitochondria following catabolism of amino acids (propionyl-CoA, crotonyl-CoA, glutaryl-CoA), fatty acids (propionyl-CoA, crotonyl-CoA, butyryl-CoA, β-hydroxybutyryl-CoA) and glucose, or the downstream tricaboxylic acid cycle (succinyl-CoA). As with acetyl-CoA, to be used by HATs, these acyl-CoAs must be exported from the mitochondria to the cytosol/nucleosol. The mechanisms underlying the export of these acyl-CoAs from the mitochondrial matrix need to be characterized, but they could rely on the carnitine shuttle system, in particular on the ability of carnitine acetyltransferase (CrAT) to generate a series of acyl-carnitines from short- and medium-chain acyl-CoAs [12]. In support of this model, CrAT-dependent generation of extra-mitochondrial acetyl-CoA [13] was reported.

Other acyl-CoAs can be generated directly in the cytosol, such as malonyl-CoA. In addition, short-chain fatty acids (SCFAs) that can be used to generate cytosolic acyl-CoAs can also be imported into cells from the extracellular environment. For example, depending on the diet or metabolic activities of intestinal microbiota, SCFAs such as propionate, succinate, butyrate or β-hydroxybutyrate (produced following fasting or a ketogenic diet), can be massively imported into the cells (for a review of the different pathways for generating cellular acyl-CoA, see [14]). Once these SCFAs have been imported into the cells, it is assumed that at least a fraction could be used by acyl-CoA synthetases to generate the corresponding SCFA-CoAs.

The first attempt to test this hypothesis was reported in 2015 on the basis of a siRNA-mediated knockdown of the nucleocytosolic acetyl-CoA synthetase, ACSS2. A decrease in the cellular concentration of ACSS2 resulted in a decreased histone crotonylation [11]. On the basis of these experimental data, the authors of this work proposed a direct generation of crotonyl-CoA by ACSS2. Following this publication, several reviews on the subject highlighted ACSS2 as an important source of crotonyl-CoA, mediating histone crotonylation (e.g. [14, 15]). Other studies have concluded that ACSS2 plays a role in promoting histone crotonylation, which mediates the reactivation of latent HIV [16]. More recently, a publicly available preprint proposes the occurrence of ACSS2-mediated H3K9 crotonylation in tubular epithelial cells [17]. However, none of the published studies directly tested the ability of ACSS2 to generate crotonyl-CoA from crotonate. In a similar manner, cytosolic ACSS has been proposed as the source of nucleocytosolic propionyl- and butyryl-CoA [14].

Here, using *in vitro* and *in vivo* experiments and structural modelling, we directly tested the ability of ACSS2 to generate SCFA-CoAs from the corresponding 3- and 4-carbon SCFAs. Our data clearly demonstrate that while ACSS2 efficiently catalyzes the production of acetyl-CoA from acetate, it is unable to generate SCFA-CoAs from 4-carbon SCFAs such as butyrate and crotonate, and has a very low activity with the 3-carbon propionate. In addition, structural considerations rationalize the inability of ACSS2 to generate acyl-CoA with a 4-carbon acyl group.

Our data therefore call for a re-examination of the idea, which has now become an established ‘fact’, of the role of ACSS2 in the generation of various SCFA-CoAs in addition to acetyl-CoA.

## Methods

### Cell culture and ACSS2 knock-down

HepG2 cells (ATCC HB-8065TM, Lot 7001058) were cultured in DMEM (Gibco, UK) and 10% FBS at 37°C and 5% CO_2_, using 6 well plates at an initial density of 3·10^5^ cells/well. After 24h, cells were transfected with siRNAs using Lipofectamine RNAiMAX (Invitrogen, MA, USA) according to the supplier’s instructions, adding per well 4.5 μL Lipofectamine and 50 pmol siRNA (On-TARGET plus™ Control pool, D-001810-10-05; or On-TARGET plus™ SMART pool ACSS2, L-010396-00-0005; both Horizon, UK). At day 2, cultures were supplemented with 1% Penicillin-Streptomycin. ACSS2 expression was verified by SDS-PAGE-immunoblotting 72 h post-transfection in whole cell lysates prepared with radioimmunoprecipitation assay buffer (RIPA). Cellular levels of CoA and acetyl-CoA were determined 96 h post-transfection by LC-MS/MS as published recently [18].

### Histone extraction and immunoblotting

Acid extraction from isolated nuclei for histone isolation was carried out 96 h post-transfection at 4°C as follows. For cell membrane lysis, adherent cells were rinsed twice with 1 mL/well ice-cold Dulbecco′s Phosphate Buffered Saline (DPBS), followed by addition of 300 μL/well of Triton extraction buffer (TEB): 0.5% Triton X-100 in DPBS, supplemented with protease inhibitor mix (complete tablets EDTA free; Roche, Germany) and phosphatase inhibitor mix (Halt™ phosphatase inhibitor cocktail; ThermoFisher Scientific, MA, USA). After gentle stirring on ice for 10 min, cells were scraped and collected in TEB. Nuclei were separated by centrifugation (300 g, 10 min, 4°C), the pellet washed with TEB and recentrifuged as before. The pellet (extracted nuclei) was then resuspended in 60 μL 0.2 N HCl, incubated overnight at 4°C, and centrifuged (300 g, 10 min, 4°C). The supernatant containing histones was then neutralized with 2 N NaOH, and protein quantified by Bradford assay (BioRad, CA, USA). Histone samples (4 μg protein/well) or whole cell lysates (see above, 40 μg protein/well) were separated on precast gels (4-20%, BioRad, CA, USA) and transferred to 0.2 μm nitrocellulose membranes (Turbo transfer pack, BioRad, CA, USA) according to the supplier’s instructions. After blocking in TBS, 0.1% (v/v) Tween and 4% (w/v) skim milk powder for 1 h at RT, the following primary antibodies were used for overnight incubation at 4°C (dilutions in parentheses): monoclonal H4K5ac, H4K8ac (both at 1:2000), H4K5cr, H4K8cr and ACSS2 (all 1:1000) from Abcam (France), Pan-Actin from (at 1:1000) from Cell Signaling (MA, USA) and polyclonal H4K5bu and H4K8bu (both at 1:1000) from ThermoFisher Scientific (MA, USA). After washing and incubation with horse-radish peroxidase-coupled secondary antibodies (anti-rabbit IgG, Jackson, PA, USA, 1:10000), luminescence was revealed with Amersham ECL™ Prime reagents (Cytiva, UK) and acquired with ImageQuant™ LAS 4000 (GE Healthcare, Il, USA).

### In vitro acetyl-CoA synthetase assay and mass spectrometry

Assays were performed in a total volume of 100 μL with following composition: 25 mM Tris buffer (pH 7.5), 6 mM MgCl_*2*_, 75 mM KCl, 3.75 mM sodium ATP (Roche, Germany), 0.5 mM CoA (Sigma Aldrich, MO, USA), and 0.5 mM SCFA substrate (sodium acetate, sodium butyrate, sodium propionate, or crotonic acid; all Sigma Aldrich, Germany); all powders were solubilized in ddH_2_O. For corresponding blanks, SCFA substrates were replaced by ddH_2_O. Reactions were started by adding to the reaction mixture: 5 μg (0.02 U) purified, cytosolic acetyl-CoA synthetase from *Saccharomyces cerevisiae* (ref. A1765, Sigma Aldrich, MO, USA; mainly ACS2, UniProt P52910 RefSeq NM_001182040.1 [19]), or 10 μg human ACSS2-overexpressing HEK-293 cell lysate (ref. LY412981, Origene, MD, USA; UniProt Q9NR19, RefSeq NM_018677 [20]) USA). The latter corresponds to < 0.0001 U per test (estimated from [20]); no generation of short chain acyl-CoAs was detectable in control HEK-293 cell lysates (LY412981, Origene). Reactions were run at 37 °C for 10 min. These conditions were shown in preliminary experiments with acetate to allow for a linear detection range without saturation (Suppl. Fig. 1). The reaction was quenched and short chain acyl-CoAs extracted by addition of 400 μL acetonitrile/methanol (v/v 1:1), incubated on ice for 30 min, centrifuged (3000 g, 5 min, 4°C), and the supernatant speed-vac dried and stored at -80°C. For acyl-CoA quantification, dry pellets were solubilized and subjected to LC-MS/MS, as recently described [18]. Analysis of acyl-CoA standards yielded a detection range of several orders of magnitude, and detection limits <200 fmol (Suppl. Fig. 2).

### Structural modelling of ACS enzymes

Structural models of human ACSS2 and ACSS3 and of *A. thaliana* ACS (*At*ACS) in the adenylate-forming conformation were downloaded from the AlphaFold Protein Structure Database (https://alphafold.ebi.ac.uk/) [21] using the UniProt accession numbers Q9NR19, Q9H6R3 and B9DGD6, respectively. The pLDDT and PAE plots for these models are shown in Suppl. Fig. 3. The N- and C-terminal domains (NTD and CTD) of these models were individually aligned onto the corresponding NTD (res. 1-516) and CTD (res. 517-647) of *Salmonella enterica* ACS (*Se*ACS) [22] (PDB 2P2F) to generate the thioester-forming conformation of these enzymes. These alignments yielded RMSD values of 0.556 Å (over 408 Cα atoms) and 0.751 Å (over 113 Cα atoms) for the ACSS2 NTD and CTD (res. 1-568 and 569-701), respectively; of 0.854 Å (over 371 Cα atoms) and 1.124 Å (over 98 Cα atoms) for the ACSS3 NTD and CTD (res. 1-555 and 556-686), respectively; and of 0.531 Å (over 391 Cα atoms) and 0.756 Å (over 107 Cα atoms) for the *At*ACS NTD and CTD (res. 1-612 and 613-743), respectively. Linker residues connecting the N- and C-terminal domains were modelled based on 2P2F residues 512-518. Regarding ligands, the coordinates for acetyl-AMP were taken from the structure of *Cryptococcus neoformans* ACS bound to acetyl-AMP (PDB 7L4G; 6R9 ligand) and aligned with the AMP ligand of PDB 2P2F. Crotonate coordinates were extracted from the structure of the crotonate-bound proton gated ion channel GLIC from *Gloeobacter violaceus* (PDB 6HJI; BEO ligand) and crotonyl-AMP was modeled by aligning the crotonate carboxylate and α methylene group atoms with the O3P, C1P and C2P atoms of the *Se*ACS propyl-AMP ligand from chain A of PDB 1PG4.

## Results

### ACSS2 knock-down does not reduce butyrylation and crotonylation at H4K5 and H4K8

Since SCFAs and acyl-CoAs play an important role in metabolically highly active liver [23], we examined the role of ACSS2 in liver-derived HepG2 cells. We induced an ACSS2-specific knock-down (KD) by siRNA (Fig. 1A) which reduced cellular ACSS2 protein levels by about 90% (Fig. 1B) and cellular acetyl-CoA levels by about 40% (Fig. 1C). Under these conditions, we analyzed acylation of two histone sites, H4K5 and H4K8, which we [10, 24, 25] and others [26, 27] have extensively studied in the past. Specific antibodies against acetylation, butyrylation and crotonylation of these sites revealed that ACSS2-KD in HepG2 cells reduced acetylation, in particular at the H4K5 site, but did not significantly change the level of butyrylation or crotonylation (Fig. 1D,E).

**Figure 1:**
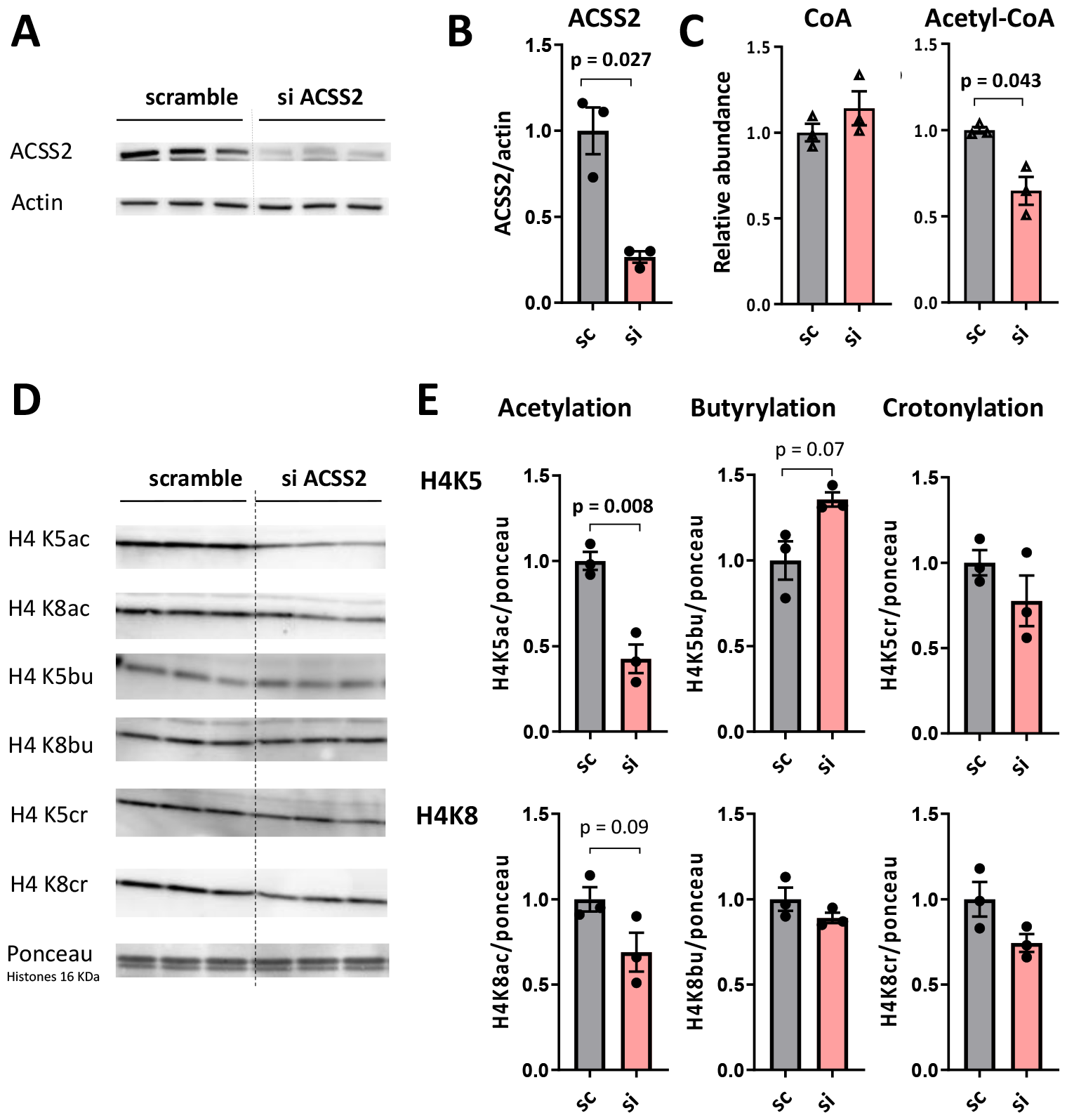
Effect of ACSS2 knock-down on the acylation status of histone 4. HepG2 cells were transfected with scramble siRNA mix or ACSS2 siRNA mix and grown for 96 h before extraction (80-90% confluence). (A) Immunoblot for siRNA efficiency (72h after transfection); actin is shown as loading control. (B) Immunoblot quantification of (A) normalized to scramble. (C) LC-MS/MS quantification of relative CoA and acetyl-CoA levels normalized to scramble. (D) Immunoblot for acetylated, butyrylated or crotonylated H4K5 and H4K8; Ponceau stain of histones (at 16 kDa) is shown as loading control. (E) Immunoblot quantification of (D) normalized to scramble. All data given as mean +/-SEM (*n*=3 independent HepG2 cultures; for MS data, each point is an average of 2 technical replicates); statistical comparison with unpaired Student’s *t*-test; values for *p* < 0.1 are given. Sc, scramble siRNA; si-ACSS2, ACSS2-specific siRNA.

### ACSS2 does not accept butyrate and crotonate as substrates in vitro

Our findings on histone acylation in the HepG2 ACSS2-KD was surprising, since an earlier study reported a strongly decreased crotonylation in HeLa cells under conditions of ACSS2-KD, using the H3K18 site as a readout [11]. Since such discrepant outcomes could be linked to the different cell lines and histone sites analyzed in the two studies, we decided to directly examine the substrate specificity of nucleocytosolic acetyl-CoA synthetases *in vitro*. Purified enzyme from the yeast *Saccharomyces cerevisiae* (mainly ACS2, homologue to mammalian ACSS2) and recombinantly expressed human ACSS2 were used in an *in vitro* assay run with supra-physiological concentrations of four SCFAs, namely acetate, propionate, butyrate, and crotonate (0.5 mM each). Reaction conditions were optimized for enzyme quantity not becoming rate-limiting with acetate as the substrate, and analysis by LC-MS/MS ensured for all acyl-CoAs of interest a linear, high-sensitivity detection. The data obtained for both yeast ACS2 and human ACSS2 were consistent (Table 1). They revealed the absence of any detectable butyryl-or crotonyl-CoA when using butyrate or crotonate as a substrate, respectively. Traces of propionyl-CoA were detectable with propionate as a substrate in a range from 0.02% (yeast) to 2.4% (human) relative to acetyl-CoA.

**Table 1:**
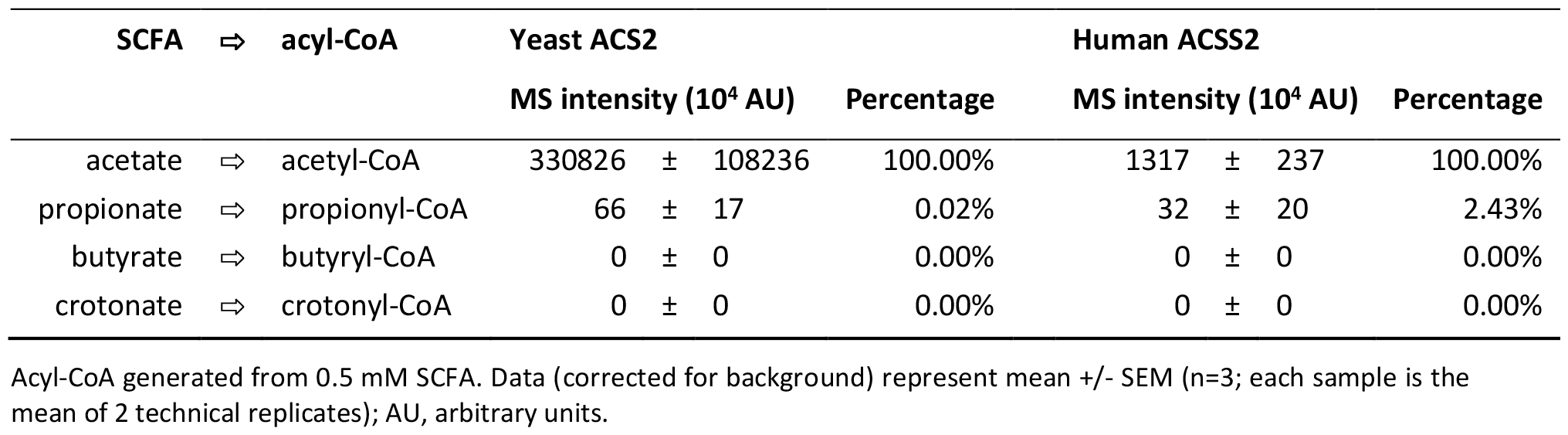
ACSS2 substrate specificity analyzed by *in vitro* assays with LC-MS/MS.

### Modeling the ACSS2 active site suggests steric hindrance for butyrl- and crotonyl-AMP intermediates

Our *in vitro* results suggest that the active site of ACSS2 may exclude SCFAs with more than three carbons as in propionate. We thus scrutinized available structural data on acetyl-CoA synthetases to verify this hypothesis. ACSS2 belongs to the class I adenylate-forming enzymes, which include aryl- and acyl-CoA synthetases, fatty acid-AMP ligases (FAAL), firefly luciferases and the adenylation (A) domain of nonribosomal peptide synthetases (NRPS) (reviewed in [28]). Crystal structures have revealed that acyl-CoA synthetases (ACSs) consist of a large (>500 residue) N-terminal domain and a small (∼120 residue) C-terminal domain, with the active site located at their interface [22, 29, 30]. These enzymes catalyse the activation of acylate to acyl-CoA in two steps. In the first (adenylation) step, the acylate substrate condenses with ATP, resulting in the release of pyrophosphate and formation of a highly reactive acyl-AMP (acyl adenylate) species (Fig. 2A). In the second (thioester formation) step, the acyl-AMP intermediate undergoes nucleophilic attack by the phosphopantothenate thiol group of CoA to yield the acyl-CoA product. These steps are catalyzed by two distinct enzyme conformations, characterized by a large relative reorientation of the N- and C-terminal domains. Although crystal structures of class I adenylate-forming enzymes are known in both conformations, the observed substrate specificity of ACSS2 can be rationalized more easily in the thioester-forming conformation because, to our knowledge, ACS structures in complex with an acyl-AMP or acylate ligand, which facilitate modeling of different sized acyl groups in the active site, are only available in this conformation.

**Figure 2.**
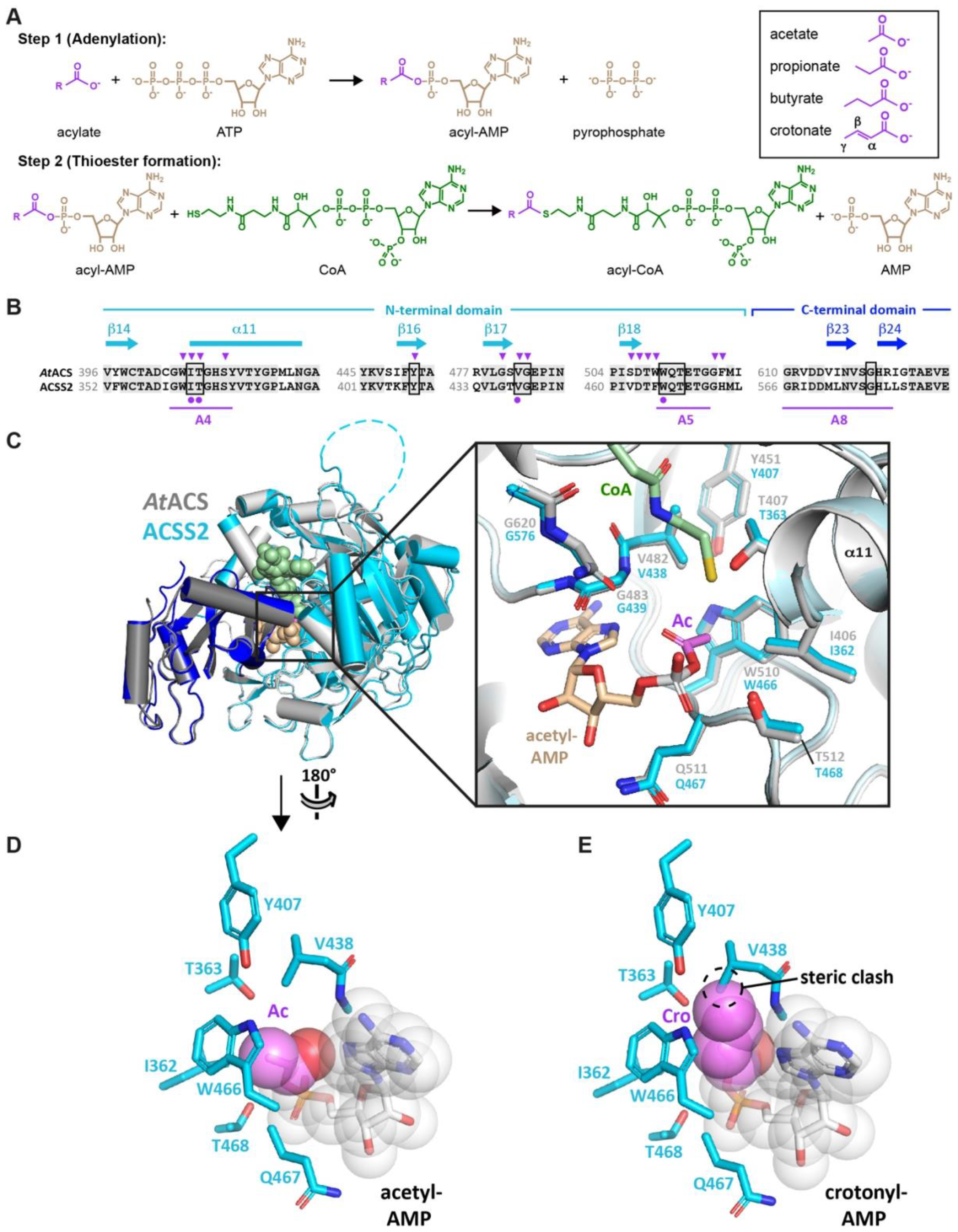
The predicted structure of the ACCS2 active site is poorly compatible with crotonyl-AMP. (**A**) Two-step reaction catalyzed by acyl-CoA synthetases. (**B**) Sequence alignment of *A. thaliana* ACS (*At*ACS) and human ACSS2 residues spanning the active site of the thioester-forming conformation. Secondary structure elements (cyan) and conserved motifs A4, A5 and A8 (magenta) of the AMP-forming family of acyl-CoA synthetases [32] are indicated. Residues implicated in substrate specificity [33] or proposed to form the carboxylate binding pocket [31] are indicated by arrowheads and circles, respectively. Boxed positions indicate residues predicted by structural modeling to lie within 6 Å of the acetyl group of acetyl-AMP. (**C**) Alignment of *At*ACS and human ACSS2 structural models. N- and C-terminal domains are shown in light and dark grey for *At*ACS and in cyan and blue for ACSS2, respectively. The structures were obtained by aligning the AlphaFold models for the N- and C-terminal domains onto the corresponding domains of *Salmonella enterica* ACS (*Se*ACS) in the thioester-forming conformation (PDB 1PG4). Residues with a low pLDDT score are either omitted (ACSS2 residues 1-23, AtACS residues 1-60) or shown as a dashed line (ACSS2 residues 257-270). The CoA (green) and acetyl-AMP (tan) ligands of *Se*ACS are shown as spheres. *Inset*. Close-up view of the active site showing the high degree of structural conservation between AtACS and human ACSS2. Residues boxed in (A) and the modeled CoA and acetyl-AMP ligands are indicated. (**D, E**) Predicted side chain environment of (C) acetyl-AMP and (D) crotonyl-AMP modeled in the active site of ACSS2. The acyl group carbon and oxygen atoms are in violet and red, respectively. Whereas the active site could easily accommodate an acetyl group, a bulkier crotonyl group is predicted to fit poorly because of steric constraints. A butyryl group would experience a similar steric hindrance.

A BLAST search of the human ACSS2 sequence against the Protein Data Bank (PDB) identified the ACS from *Salmonella enterica* (*Se*ACS) as the top hit, with 51.8% identity over 93% of the query sequence. The structure of *Se*ACS is known in the thioester-forming conformation bound to CoA and to either propyl-AMP or acetate and AMP [29, 30]. The ∼40-fold higher selectivity of ACSS2 for acetate versus propionate and the inactivity towards butyrate and crotonate (Table 1) is strongly reminiscent of the substrate selectivity reported for *A. thaliana* ACS (*At*ACS), whose catalytic efficiency (*k*_cat_/*K*_m_) is 55-fold higher with acetate than with propionate and which shows no activity with longer chain carboxylates [31]. ACSS2 and *At*ACS share an overall sequence identity of 51%, which increases to 78% identity over regions spanning the conserved motifs defining the thioester-forming active site (Fig. 2B).

Structural models of these enzymes are available from the AlphaFold Protein Structure Database (AlphaFold DB) in the adenylate-forming conformation. We therefore generated the thioester-forming conformations by individually aligning their N- and C-terminal domains onto those of *Se*ACS, and modeled the atomic coordinates of acetyl-AMP and crotonyl-AMP from available ligand-bound crystal structures, as detailed in the Methods. The resulting ACSS2 and *At*ACS models were remarkably similar, sharing an RMSD of 0.495 Å over 581 Cα atoms (Fig. 2C). Notably, all residues within contact distance of the acetyl-AMP acetyl group are identically conserved and share an all-atom RMSD of 0.236 Å over 72 atoms, consistent with the similar substrate selectivities of these two enzymes. Furthermore, whereas all atoms of the acetyl-AMP ligand are well accommodated by the surrounding side chains in our ACSS2 model (Fig. 2D), replacing the acetyl group by a bulkier crotonyl group would result in a steric clash between the crotonyl γ-methyl group and the Val438 side chain (Fig. 2E). Likewise, the presence of a γ-methyl group in butyryl-AMP and its absence in propionyl-AMP (Fig. 2A) are consistent with the ability of ACSS2 to accept propionate, but not butyrate, as a substrate.

Thus, structural modeling of the ACSS2 active site rationalizes the substrate selectivity observed for this enzyme.

### The predicted structure of the ACSS3 active site is compatible with butyryl- and crotonyl-AMP

We next wondered whether the active site of ACSS3, a mitochondrial paralogue of ACSS2 (32% sequence identity), might be compatible with butyryl- and crotonyl-AMP, given that the amino acid sequence of ACSS3 differs significantly from that of ACSS2 and *At*ACS within regions spanning the thioester-forming active site (Suppl. Fig. 4A). As described above for ACSS2 and *At*ACS, we modeled ACSS3 in the thioester-forming conformation using the ACSS3 model from the AlphaFold DB and the crystal structure of *Se*ACS. The ACSS2 and ACSS3 active sites differ most notably at four positions: ACSS2 residues Ile362, Tyr407 and Val438 are replaced by the smaller ACSS3 residues Val345, Phe391 and Ala424, respectively, while the polar residue Thr363 is replaced by the hydrophobic residue Val346 (Suppl. Fig 4B). These substitutions are predicted to create an additional empty space above the acetyl-AMP acetyl group surrounded by a hydrophobic environment (Suppl. Fig. 4C). Modelling shows that a crotonyl or butyryl group could easily be accommodated within this space without steric hindrance (Suppl. Fig. 4D). This finding suggests mitochondrial ACSS3 as a plausible candidate for generating butyrl- and crotonyl-CoA from butyrate and crotonate, respectively.

## Discussion

Direct *in vitro* measurement of SCFA-CoA synthesis mediated by cytosolic human and yeast acetyl-CoA synthetases (ACSS2, ACS2), reported here, clearly demonstrated the inability of these enzymes to synthesize longer-chain acyl-CoA, whereas, as expected, they efficiently produce acetyl-CoA from acetate. We also note that ACSS knockdown does not significantly change H4K5 and H4K8 butyrylation and crotonylation in HepG2 cells. However, focusing on crotonyl-CoA, we observe in the literature that the idea of ACSS2 ensuring nucleocytosolic production of crotonyl-CoA is widespread and is even considered an established ‘fact’. However, this belief seems to be based solely on the reported effect of ACSS2 knockdown on histone crotonylation [11], without any further experimental validation.

In order to explain the discrepancy between the published data and our cellular and direct *in vitro* and structural analyses, we propose that the reported effect of ACSS2 knockdown on histone crotonylation could be entirely indirect. Indeed, a decrease in cellular acetyl-CoA production, after ACSS2 knockdown would in turn affect the accumulation of fatty acids and hence β-oxidation, known as a factor that can mediate histone crotonylation [25, 34]. In addition, a decrease in ACSS2 could also specifically affect an acetylation-dependent gene expression program, which could control the production of crotonyl-CoA, through different other metabolic pathways. It is also possible that these ACSS2 and acetylation-dependent genes encode the yet unknown SCFA-synthetase responsible for crotonyl-CoA production. Indeed, recent elegant studies have shown that yeast ACS2 mediates a redistribution of histone acetylation across the genome and therefore controls a subsequent change in gene expression in response to a specific cell signaling system [35, 36].

However, the literature clearly shows that cells are able to use extracellular crotonate to ensure histone crotonylation. At the physiological level, it has been shown that the intestinal microbiota contribute to increased histone crotonylation in the cryptic fraction of the intestinal epithelium of the small intestine and colon [37]. In addition, supplementation of cell culture media with crotonate has been shown to increase histone crotonylation [11], suggesting the ability of cells to import crotonate from their environment and generate crotonyl-CoA that can be used by HATs to crotonylate histones (see e.g. [37]).

It is therefore reasonable to assume that one or more cellular acyl-CoA synthetases could be responsible for crotonyl-CoA generation. However, our data show that, contrary to the widely accepted idea, this enzyme is not ACSS2. This conclusion also agrees with earlier data showing that SCFA-CoA synthetases from yeast and mammals, although active towards acetate, can hardly use butyrate as a substrate [38].

The remaining question is therefore which enzyme(s) is(are) capable of using SCFAs other than acetate for the synthesis of short-chain acyl-CoAs, including crotonyl-CoA. The human genome encodes 26 distinct acyl-CoA synthetases, which may be involved in fatty acid activation depending on the length of the acyl chain, although there could be a significant overlap in chain length specificity [39]. We found that the active site of at least one of these, ACSS3, is predicted to be sterically compatible with a crotonyl-AMP intermediate. Our structural study and literature review highlighted the stereochemical requirements for the use of SCFAs longer than propionate and showed why ACSS2 is not an eligible candidate for the synthesis of crotonyl-CoA.

## Supplemental material

**Supplemental Figure 1:**
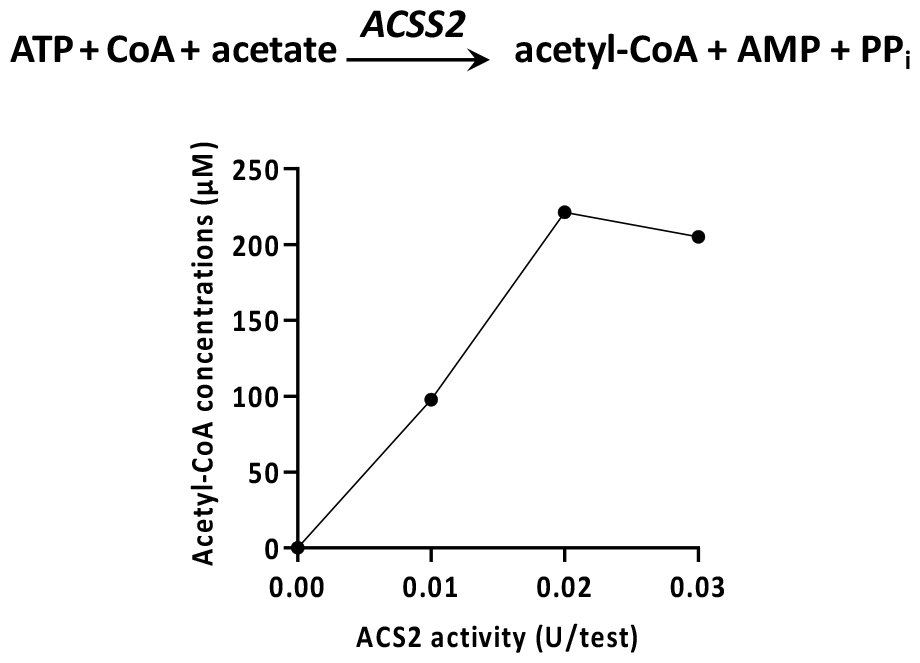
In vitro acetyl-CoA synthetase assay. Preliminary experiment to check assay linearity and saturation, using yeast ACS2 and acetate as substrate. The reaction mix (100 μL) was quenched and ACS2 enzyme was precipitated by adding ice-cold 0.5 N perchloride acid (50 μL) and incubation on ice for 30 min. After centrifugation (3000 g, 12 min, 4°C), the supernatant containing acetyl-CoA product was neutralized with 5 M K_2_CO_3_ and centrifuged again (3000 g, 12 min, 4°C). Acetyl-CoA in the supernatant was quantified by HPLC [18].

**Supplemental Figure 2:**
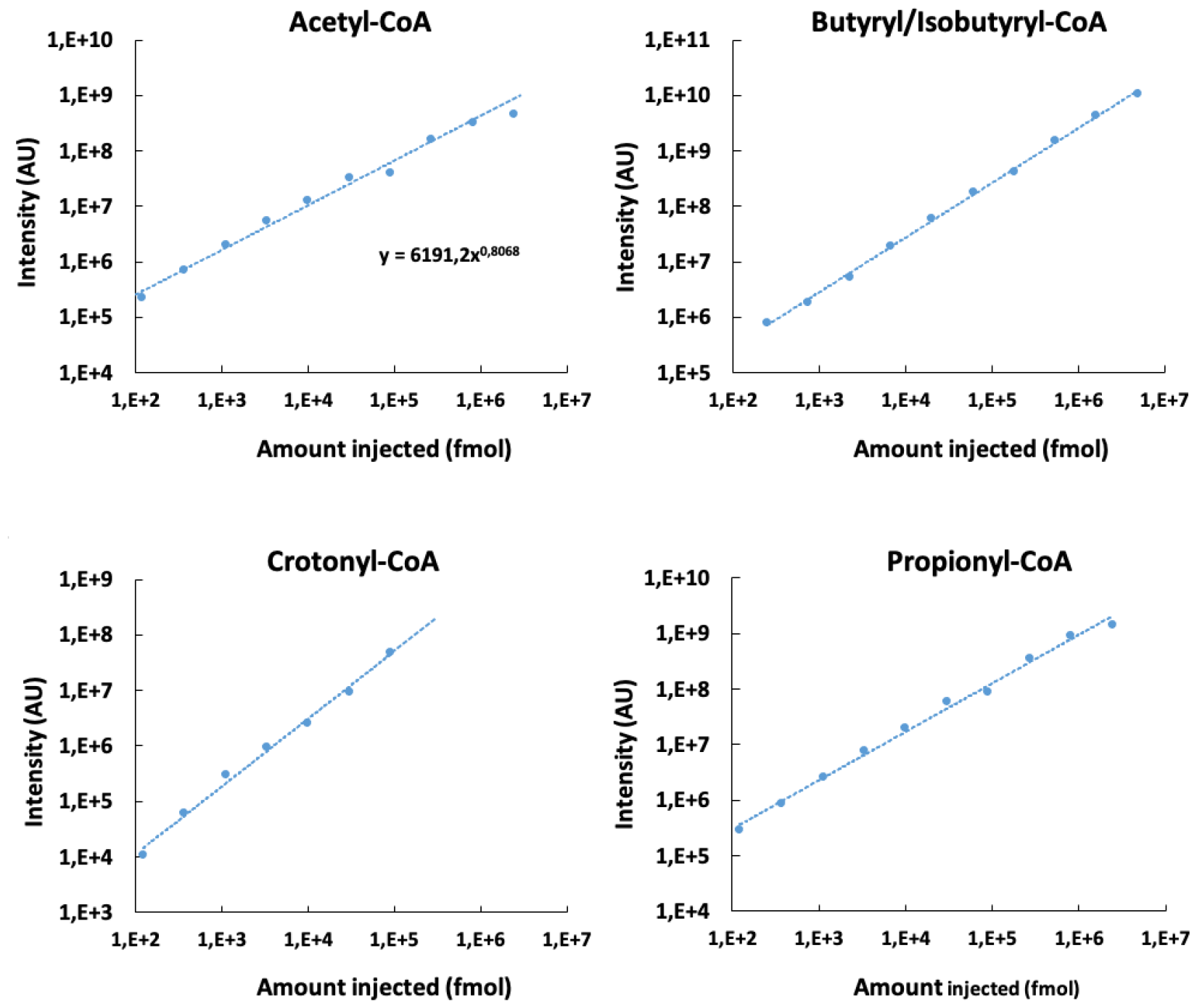
LC-MS/MS signal as a function of injected acyl-CoA standard. A dilution series of a mixture of acyl-CoA standards was quantified by LC-MS/MS as published recently [18]. An exponential regression line is indicated. Note that data are presented with a double logarithmic scale.

**Supplemental Figure 3.**
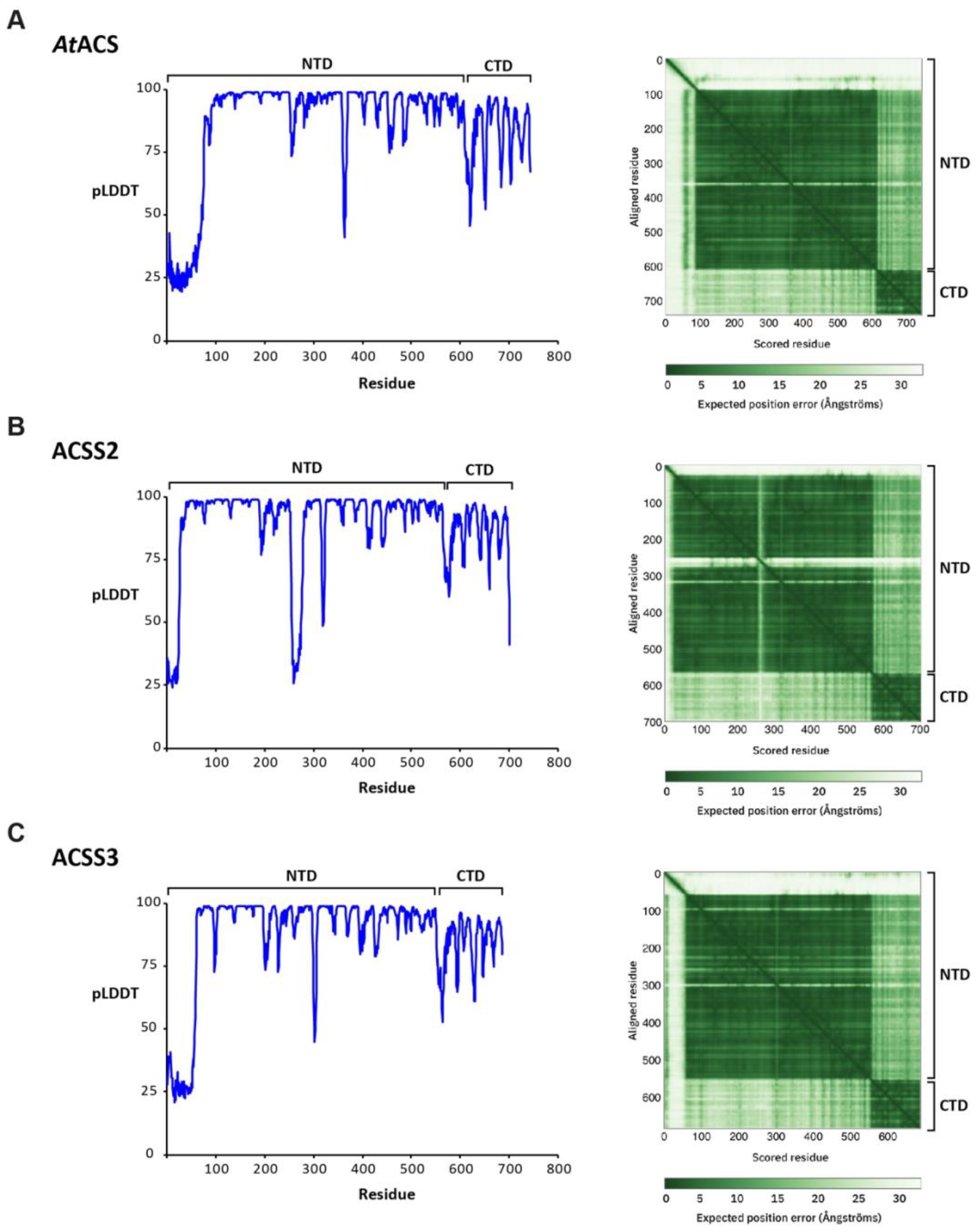
Confidence plots for AlphaFold models of ACS enzymes. Plots of pLDDT score (left) and predicted aligned error (PAE) are shown for (A) *At*ACS, (B) human ACSS2 and (C) human ACSS3. The N-terminal domain (NTD) and C-terminal domain (CTD) limits are indicated.

**Supplemental Figure 4.**
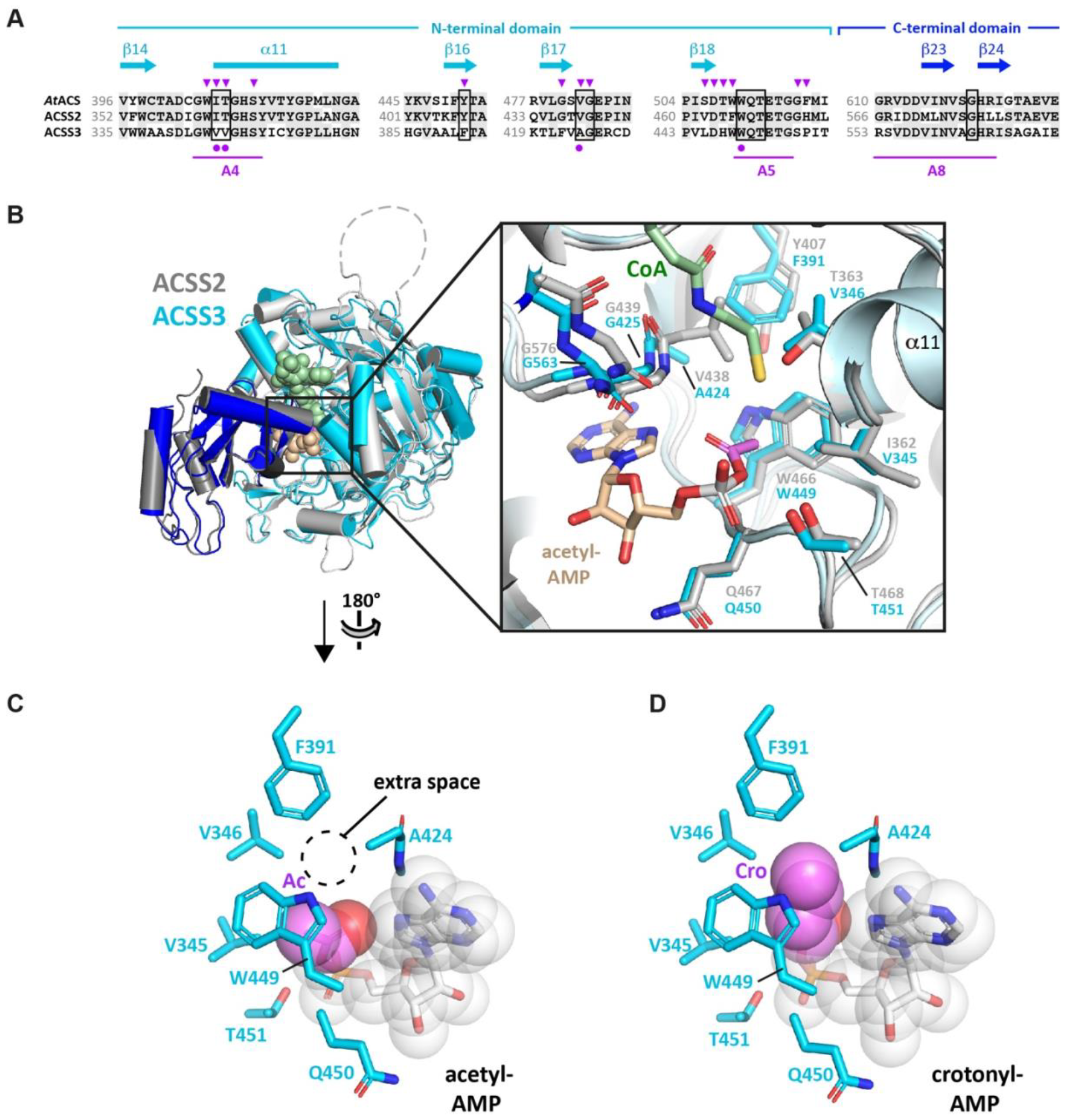
The predicted structure of the ACCS3 active site can feasibly accommodate butyryl- and crotonyl-AMP. (**A**) Sequence alignment of *A. thaliana* ACS (*At*ACS) and human ACSS2 and ACSS3 residues spanning the active site. Secondary structure elements (cyan) and conserved motifs A4, A5 and A8 (magenta) of the AMP-forming family of acyl-CoA synthetases [32] are indicated. Residues implicated in substrate specificity [33] or proposed to form the carboxylate binding pocket [31] are indicated by arrowheads and circles, respectively. Boxed positions indicate residues predicted by structural modeling to lie within 6 Å of the acetyl group of acetyl-AMP. (**B**) Alignment of human ACSS2 and ACSS3 structural models. N- and C-terminal domains are shown in light and dark grey for ACSS2 and in cyan and blue for ACSS3, respectively. The structures were obtained by aligning the AlphaFold models for the N- and C-terminal domains onto the corresponding domains of *Salmonella enterica* ACS in the thioester-forming conformation (PDB 1PG4). *Inset*. Close-up view of the active site showing notable structural differences between ACSS2 and ACSS3. Residues boxed in (A) and the modeled CoA and acetyl-AMP ligands are indicated. (**C, D**) Predicted side chain environment of (C) an acetyl-AMP and (D) a crotonyl-AMP modeled in the active site of ACSS3. The acyl group carbon and oxygen atoms are in violet and red, respectively. The additional space next to the acetyl group is predicted to allow a bulkier butyryl or crotonyl group to fit comfortably within the active site.

